# The impact of identity-by-descent on fitness and disease in natural and domesticated *Canid* populations

**DOI:** 10.1101/2020.11.16.385443

**Authors:** Jazlyn A. Mooney, Abigail Yohannes, Kirk E. Lohmueller

## Abstract

Domestic dogs have experienced population bottlenecks, recent inbreeding, and strong artificial selection. These processes have simplified the genetic architecture of complex traits, allowed deleterious variation to persist, and increased both identity-by-descent (IBD) segments and runs of homozygosity (ROH). As such, dogs provide an excellent model for examining how these evolutionary processes influence disease. We assembled a dataset containing 4,414 breed dogs, 327 village dogs, and 380 wolves genotyped at 117,288 markers and phenotype data for clinical and morphological phenotypes. Breed dogs have an enrichment of IBD and ROH, relative to both village dogs and wolves and we use these patterns to show that breed dogs have experienced differing severities of bottlenecks in their recent past. We then found that ROH burden is associated with phenotypes in breed dogs, such as lymphoma. We next test the prediction that breeds with greater ROH have more disease alleles reported in Online Mendelian Inheritance in Animals (OMIA). Surprisingly, the number of causal variants identified correlates with the popularity of that breed rather than the ROH or IBD burden, suggesting an ascertainment bias in OMIA. Lastly, we use the distribution of ROH across the genome to identify genes with depletions of ROH as potential hotspots for inbreeding depression and find multiple exons where ROH are never observed. Our results suggest that inbreeding has played a large role in shaping genetic and phenotypic variation in dogs, and that there remains an excess of understudied breeds that can reveal new disease-causing variation.

**Significance Statement:** Dogs and humans have coexisted together for thousands of years, but it was not until the Victorian Era that humans practiced selective breeding to produce the modern standards we see today. Strong artificial selection during the breed formation period has simplified the genetic architecture of complex traits and caused an enrichment of identity-by-descent (IBD) segments in the dog genome. This study demonstrates the value of IBD segments and utilizes them to infer the recent demography of canids, predict case-control status for complex traits, locate regions of the genome potentially linked to inbreeding depression, and to identify understudied breeds where there is potential to discover new disease-associated variants.

## Introduction

Identity-by-descent (IBD) segments are stretches of the genome that are inherited from a common ancestor and that are shared between at least two genomes in a population (Figure S1). Runs of homozygosity (ROH) form when an individual inherits the same segment of their genome identically by descent from both parents. Thus, ROH can be viewed as a special case of IBD, where IBD occurs within an individual rather than shared between individuals (1) (Figure S1). Recent consanguinity generates an increase in ROH (2, 3), while decreases in population size can generate IBD (4, 5).

Dogs provide an excellent model system for testing how ROHs and IBD patterns impact complex traits and reproductive fitness. The unique demographic and selective history of dogs includes a domestication bottleneck coupled with subsequent breed formation bottlenecks and strong artificial selection. This demography has allowed the persistence of deleterious variation, simplified genetic architecture of complex traits, and an increase in both ROH and IBD segments within breeds (6–11). Specifically, the average F_ROH_ was approximately 0.3 in dogs (12), compared to 0.005 in humans, computed from 1000 Genomes populations (13). The large amount of the genome in ROHs in dogs, combined with a wealth of genetic variation and phenotypic data (7, 10, 12, 14–16) allow us to test how these factors influence complex traits. Further, many of the deleterious alleles within dogs likely arose relatively recently within a breed, and dogs tend to share similar disease pathways and genes with humans (9, 17, 18), making our results relevant for complex traits in humans.

Despite IBD segments and ROHs being ubiquitous in genomes, the extent to which they affect the architecture of complex traits as well as reproductive fitness has remained elusive. Given that ROH are formed by inheritance of the same ancestral chromosome from both parents, there is a much higher probability of the individual to become homozygous for a deleterious recessive variant (13, 19) that was once carried in a heterozygous form, leading to a reduction in fitness. This prediction was verified in recent work in non-human mammals that has shown that populations suffering from inbreeding depression tend to have an increase in ROHs (20, 21). ROHs in human populations are enriched for deleterious variants. However the extent to which these impact phenotypes has not been demonstrated (13, 19, 22). Along these lines, several studies have associated an increase in ROHs with complex traits in humans (23–28), though some associations remain controversial (29–33). Determining how ROHs and IBD influence complex traits and fitness could provide a mechanism for differences in complex trait architecture across populations that vary in their burden of IBD and ROH.

Here, we use IBD segments and ROH from 4,741 breed dogs and 379 wolves to determine the recent demographic history of dogs and wolves and establish a connection between recent inbreeding and deleterious variation associated with both disease and inbreeding depression. This comprehensive dataset contains genotype data from 172 breeds of dog, village dogs from 30 countries, and gray wolves from British Colombia, North America, and Europe. We use IBD segments to infer the recent demographic history of these canids. In agreement with previous studies, we find that breed dogs experienced breed formation bottlenecks of varying degrees, and as expected, find no evidence of a breed formation bottleneck in village dogs or wolves (7). We test for an association with the burden of ROH and case-control status for a variety of complex traits. We find that an increase in ROH is associated with lymphoma, portosystemic vascular anomalies, and cranial cruciate ligament disease within breed dogs. Remarkably, we also find that the number of disease-associated causal variants identified in a breed is positively correlated with breed popularity rather than burden of IBD or ROH in the genome, suggesting ascertainment biases also exist in databases of dog disease mutations. Lastly, we identify multiple loci that may be associated with inbreeding depression by examining localized depletions of ROH across dog genomes.

## Results

### Global patterns of genetic diversity across dogs and wolves

To examine genetic diversity in dogs and wolves, we merged three previously published genotype array-based datasets (14–16). As an initial quality check, we used principal component analysis (PCA) to examine the relationship between domesticated dogs, village dogs, and wolves (Figure1A). We observed a split between dogs and wolves on the first PC. The dogs that fall closest to wolves trace their origins back to Australasia (Figure 1A), which has been previously shown to be the origin of some of the more ancient dog breeds (10, 34). When we performed a PCA with only breed dogs, they clustered by clade (Figure S2). Clades are composed of multiple breeds defined in previous work (7, 30). We also observed separation based on the geographic location of wolf populations, which originate from populations from Europe or North America. A PCA with only wolves showed clear clustering by the location of where samples originated from, which includes the United States, Mexico, or Europe (Figure S3).

**Figure 1.**
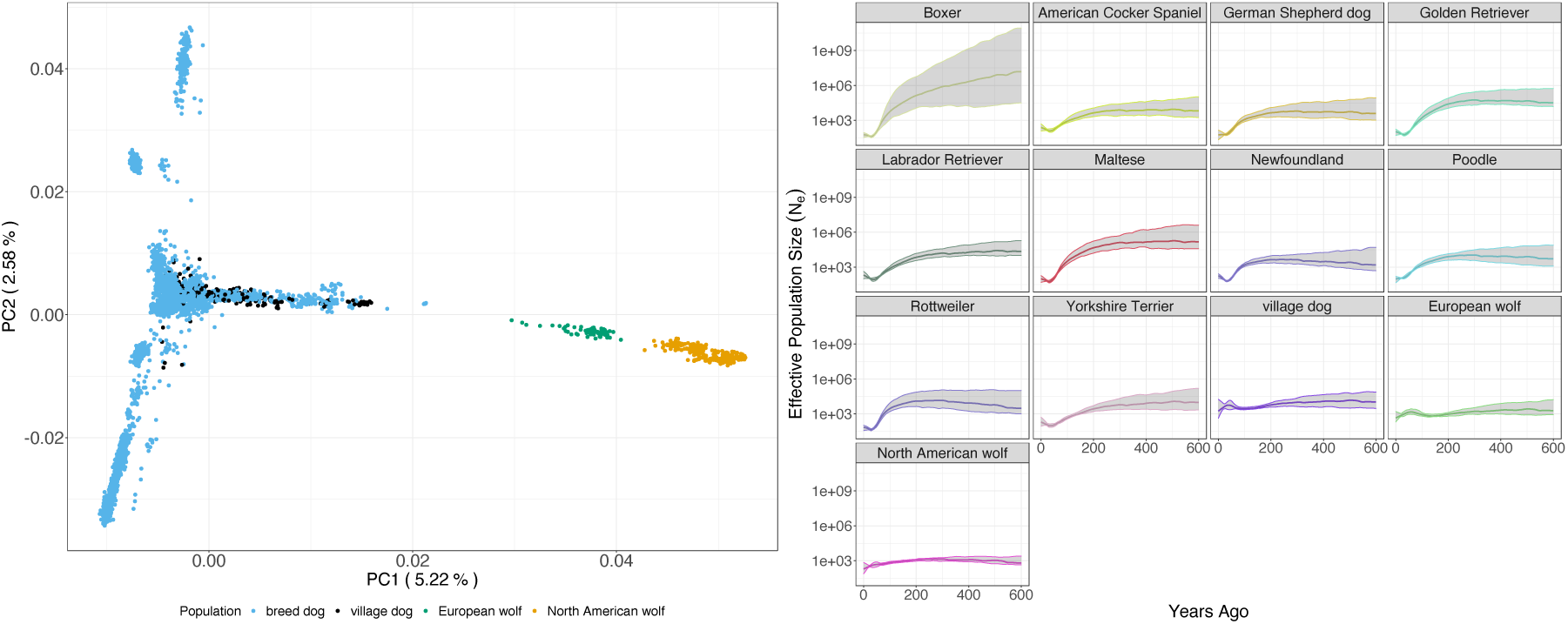
**A)** PCA of breed dogs, village dogs, and wolves. The 350 village dogs were sampled from 32 different countries around the world and the 379 wolves were sampled from populations across Europe and North America. **B)** Effective population size (N_e_) trajectories through time of breed dogs, village dogs, and wolves were inferred using IBDNe, when there were at least 50 unrelated samples per population (see *Methods*). Shaded regions in the plots indicate the 95% confidence interval of the inferred population size at each time point.

### Breed-specific bottlenecks are captured in IBD patterns

We next turned our attention to IBD patterns and called IBD segments using IBDSeq (35). We then tested whether the recent demographic history of dogs, consisting of breed formation bottlenecks within the last few hundred years (11, 18, 36), had an impact on the IBD patterns. Although the demographic history of dogs has been well studied over the years, the majority of these works have focused on the origin of dogs thousands of years ago, and their geographic origin remains an ongoing point of contention (10, 34, 37–40). We inferred the recent demographic history of 10 standard breeds of dog, village dogs, and gray wolves from North America and Europe (Figure 1B) using patterns of IBD sharing between individuals (41). When conducting demographic inferences using IBDNe, we restricted Mour analyses to populations with at least 50 individuals (see Discussion). The IBDNe analyses show that all breed dogs have experienced a domestication bottleneck followed by another severe bottleneck approximately 200 years ago which corresponds with modern breed formation during the Victorian Era (1800s) (Figure 1B). Though the strength of breed formation bottleneck varied across breeds, and was less pronounced in mixed breeds, all bottlenecks were followed by a subsequent increase in population size. Notably, the Maltese and Rottweiler appeared to have undergone the most severe bottlenecks with the Golden Retriever and German Shepherd dog close behind. The boxer also experienced a severe bottleneck, but this bottleneck may be linked to reference bias (CanFam3.1 reference genome is from a boxer) as the confidence intervals are also largest for the boxer. Village dogs (feral street dogs) and gray wolves do not appear to have experienced the domestication bottleneck which is concordant with their history (7). Instead, the village dogs showed a much weaker prolonged bottleneck followed by an increase in population size (Figure 1B). The European gray wolves have a demographic trajectory similar to the village dogs, and the American gray wolves appear to have just experienced a prolonged decline in population size (Figure 1B), consistent with recent ecological studies (14, 42, 43). Thus, the formation of dog breeds in the last 200 years has increased the amount of IBD within dogs compared to wolves and village dogs.

### Long ROH are enriched in most breed dogs

We next sought to examine the burden (total amount) of long ROH (greater than 2Mb) across breeds. We observed that F_ROH_, the proportion of the genome within a long ROH, varies across breed and by extension clade (Figure S4). The majority of breed dogs contained a larger amount of their genome in ROH and thus a larger average value of F_ROH_, a likely consequence of having experienced both a domestication and breed formation bottleneck as well as inbreeding (3). The Jack-Russell Terrier was the exception and had reduced ROH relative to other breed dogs, similar to previous works where it was found to be an outlier (10, 11). In village dogs, we observed that mean values of F_ROH_ fall much closer to what we observed in the wolves. This is expected since village dogs only experienced the domestication bottleneck and were left to reproduce without selective breeding. Lastly, we examined ROH among wolves, the European gray wolf has mean values of F_ROH_ that are comparable to the village dogs and markedly lower than the American gray wolves. The American gray wolf exhibited increased mean values of F_ROH_, likely due to having experienced a recent bottleneck (Figure 1B) as a result of being pushed to near extinction (44).

### Disease traits are associated with ROH burden

We hypothesize that the prevalence of ROH and identity-by-descent (IBD) segments could be associated with recessive genetic disease in each breed (Figure S1). ROH form when an individual inherits the same segment of their genome identically by descent from both parents (1), and the formation of ROH results in an increased probability of the individual to be homozygous for a deleterious recessive variant (19, 45). Thus, we predict that breeds with large amounts of ROH and IBD segments will have an increased incidence of recessive-disease. We tested this hypothesis using data from 4,342 dogs where we had case-control status for subsets of the data across 8 clinical and morphological phenotypes (Figure 3). For the majority of traits, there was not a significant association with ROH burden, even when stratified by breed. However, we observed an excess of associations at a nominal significance level (p<0.05) compared to what was expected under the null hypothesis of no trait associations (6 observed associations vs. 1.45 associations expected under the null, p = 0.0027, binomial test; Figure 2). We observed a significant association between the burden of ROH and case-control status for five traits: portosystemic vascular anomalies (PSVA) in Yorkshire Terriers (β = -0.394 & p < 0.027), lymphoma within both Labrador (β = -0.604 & p < 0.0340) and Golden Retrievers (β = 0.913 & p < 0.001), cranial cruciate ligament disease (CLLD) in Labrador Retrievers (β = -0.403 & p < 0.003), elbow dysplasia (ED) across all breeds (β = 0.238 & p < 0.047), and mast cell tumors (MCT) across all breeds (β = 0.286 & p < 0.027). For lymphoma in Golden Retrievers, case-status is positively associated with the amount of the genome within an ROH (OR = 2.491, Table S1) and on average cases carried more ROH than controls (Figure S5). Conversely, ROH appeared to show a protective effect against developing PSVA in Yorkshire Terriers, or CLLD and lymphoma in Labrador Retrievers.

**Figure 2.**
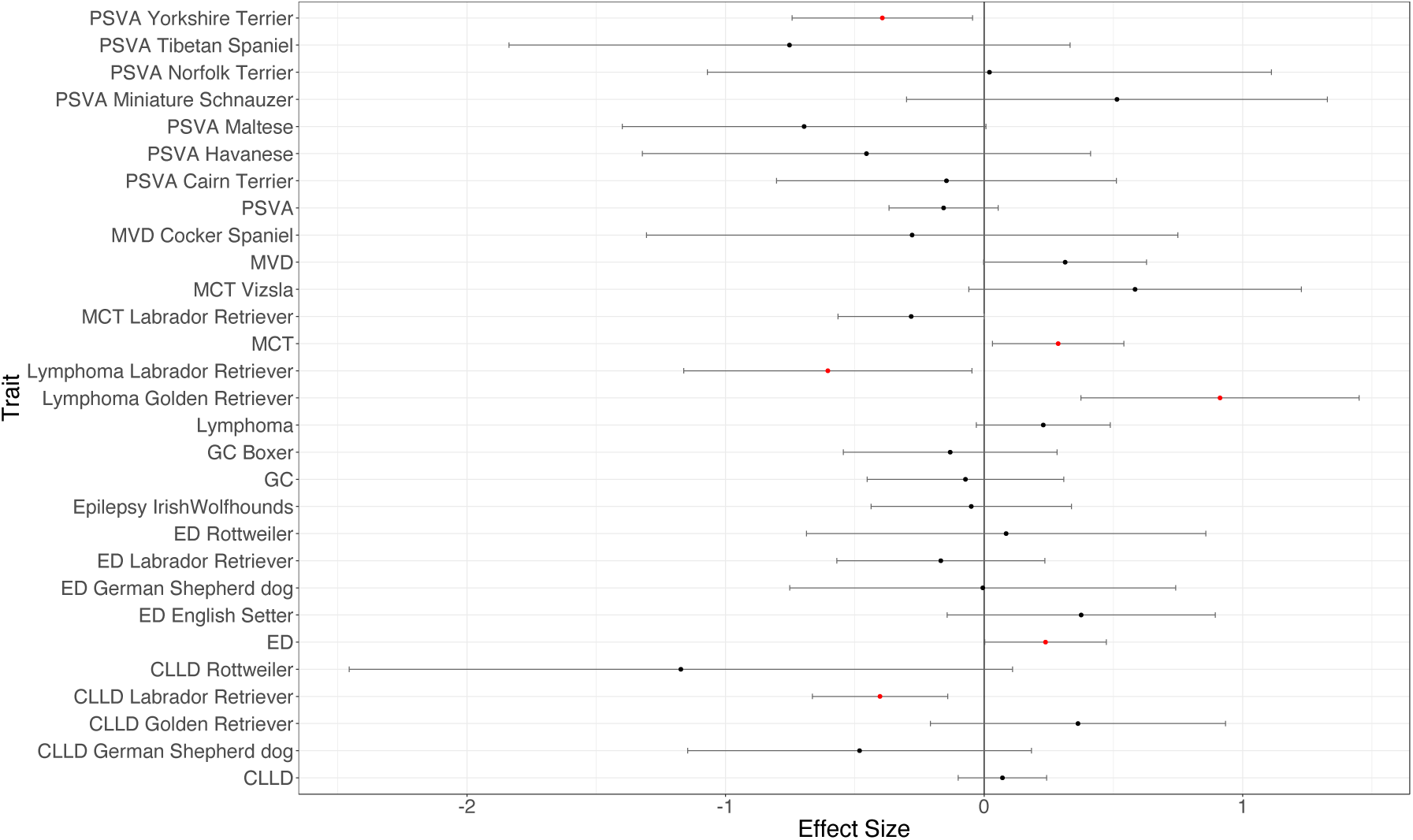
Association of ROH burden with eight quantitative traits. Results are presented both stratified by breed and across all breeds. A significant effect of ROH burden on a trait (p < 0.05) is indicated with a red point. An effect size greater than 0 indicates an increase in ROH with the trait or disease status, and less than 0 represents the converse. Phenotype abbreviations: portosystemic vascular anomalies (PSVA); mitral valve degeneration (MVD); mast cell tumor (MCT); granulomatous colitis (GC); elbow dysplasia (ED); and cranial cruciate ligament disease (CCLD). These results use shared ROH (SR) correction for populations stratification (see *Methods*). Table S1 contains sample sizes, effect sizes, odds-ratio, confidence intervals, and p-values. Figure S9 shows the uncorrected results as well as results using the genotype relatedness matrix.

**Figure 3.**
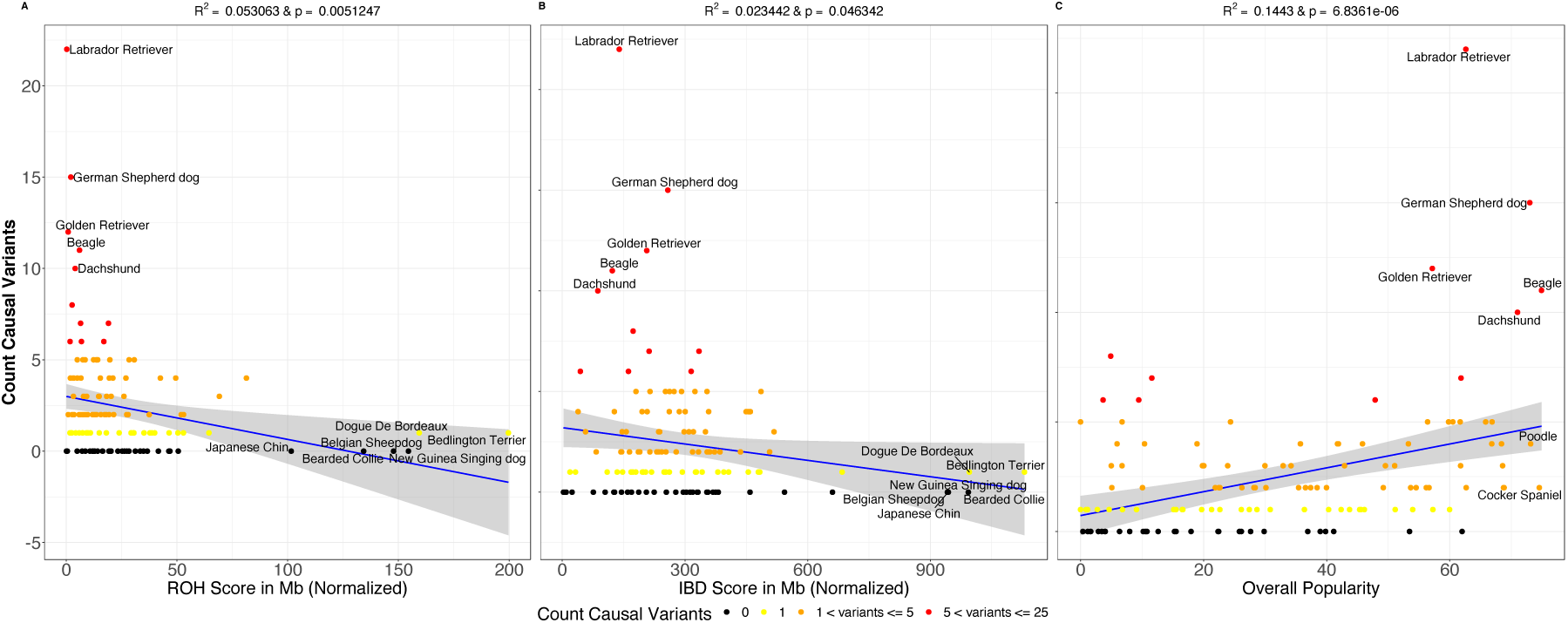
Breed popularity is positively correlated with number of causal variants identified in OMIA. Here we show the correlation between causal variants identified in each breed that have been reported in OMIA and three different metrics. The shaded regions in each plot represent the confidence interval on the regression line. **A)** The correlation between within-breed ROH and the total number of causal variants associated disease identified in the breed that have been reported in OMIA. **B)** The correlation between within-breed IBD and the total number of causal variants associated with disease identified in the breed that have been reported in OMIA. **C)** The correlation between breed popularity over time and the total number of causal variants associated with disease.

### Breed popularity, rather than ROH, correlates with Mendelian disease incidence

If ROH and IBD increase the homozygosity of disease alleles, we hypothesize that dogs with the largest amount of ROH and/or IBD would carry the most disease associated variants. We tested this hypothesis using data from Online Mendelian Inheritance in Animals (OMIA) which included a count of causal variants identified in each breed. We observed that breeds with the lowest amounts of ROH/IBD have the most identified causal variants (Figure 3 A and B). Further, those breeds with the most ROH/IBD have no causal variants identified, apart from the Kerry Blue Terrier that has been used in a single study (Figure 3 A and B). This finding was unexpected, and so we sought additional factors that might explain the variation in the number of OMIA variants per breed. We chose to examine the popularity of different dog breeds over time using data compiled by the American Kennel Club (AKC). We observed a strong positive correlation between the overall breed popularity with the number of causal variants identified in each breed (R^2^ = 0.168 & p = 1.145 x 10^− 06^) (Figure 3C). We find that the most popular breeds, such as the Retrievers, have the most causal variants identified in genomic studies. Given that we do not observe a positive relationship between IBD or ROH and the total number of causal variants in a breed, and that breeds with excess amounts of IBD and/or ROH have almost no causal variants identified, our results indicate that there are large-scale ascertainment biases in OMIA reported disease associated variants in dogs. More causal variants have been identified in the more popular breeds, containing fewer ROH, rather than those breeds with an increased prevalence of the associated Mendelian disease containing more ROH. Our results suggest that there are many understudied breeds that may be valuable for variant discover such as the Bearded collie, Belgian Sheep dog, Bedlington Terrier, or Dogue de Bordeaux, and these breeds are prone to serious health conditions according to the American Kennel Club (akc.org/dog-breeds).

### ROH reveal genes with recessive lethal mutations

Given the relatively high values of F_ROH_ observed for breed dogs, much of the genome should be in in ROH in at least one of the 4,342 individuals in our study. We hypothesized the genes not contained within a ROH in any individual or showing a deficit of ROH compared to the rest of the genome, contain recessive lethal variants, because individuals homozygous for these mutations are not viable. Across 4,342 dogs, we observed 27 genes (coordinates available on <u>GitHub</u>) where at least one exon does not overlap a ROH in any individual. To test whether this is unusual, we permuted the locations of the ROHs within each individual and re-counted the number of genes with an exon not containing a ROH. We found that if ROHs were randomly distributed across the genome, we would expect to see ROH in all exons across genes (Figure S6). Thus, there are more genes not overlapping ROHs than expected by chance (p < 0.0001) suggesting the presence of segregating recessive lethal mutations across breed dogs.

To test whether these 27 genes may have a functional effect or impact fitness, we intersected these 27 genes with the 90^th^ percentile constrained coding regions (CCRs) identified in human populations (46). CCRs were found to be enriched for disease-causing variants, especially in dominant Mendelian disorders, and the 95^th^ percentile CCRs may be enriched for embryonic lethal mutations (46). We expect that genes containing recessive lethal mutations would be conserved across species. We tested whether the genes not overlapping ROH in any dogs were enriched for CCRs. On average, one would expect to see 18 of 27 genes above the 90^th^ percentile CCRs, based on random gene sets created by resampling genes and intersecting with 90^th^ percentile CCRs 100,000 times (see Methods). However, we observed that 23 out of 27 of our genes not overlapping ROH in dogs fall above the 90^th^ percentile of the CCR distribution (p = 0.025) (Figure 4). Additionally, we observed a 2.94-fold enrichment of non-ROH genes relative to ROH genes in CCRs (p = 0.041, Fisher’s Exact) (Figure 4). Taken together, these results suggest that the genes with an exon not overlapping a ROH in dogs are enriched for exons devoid of variation in humans. Thus, these genes may be targets of strongly deleterious mutations affecting viability.

**Figure 4.**
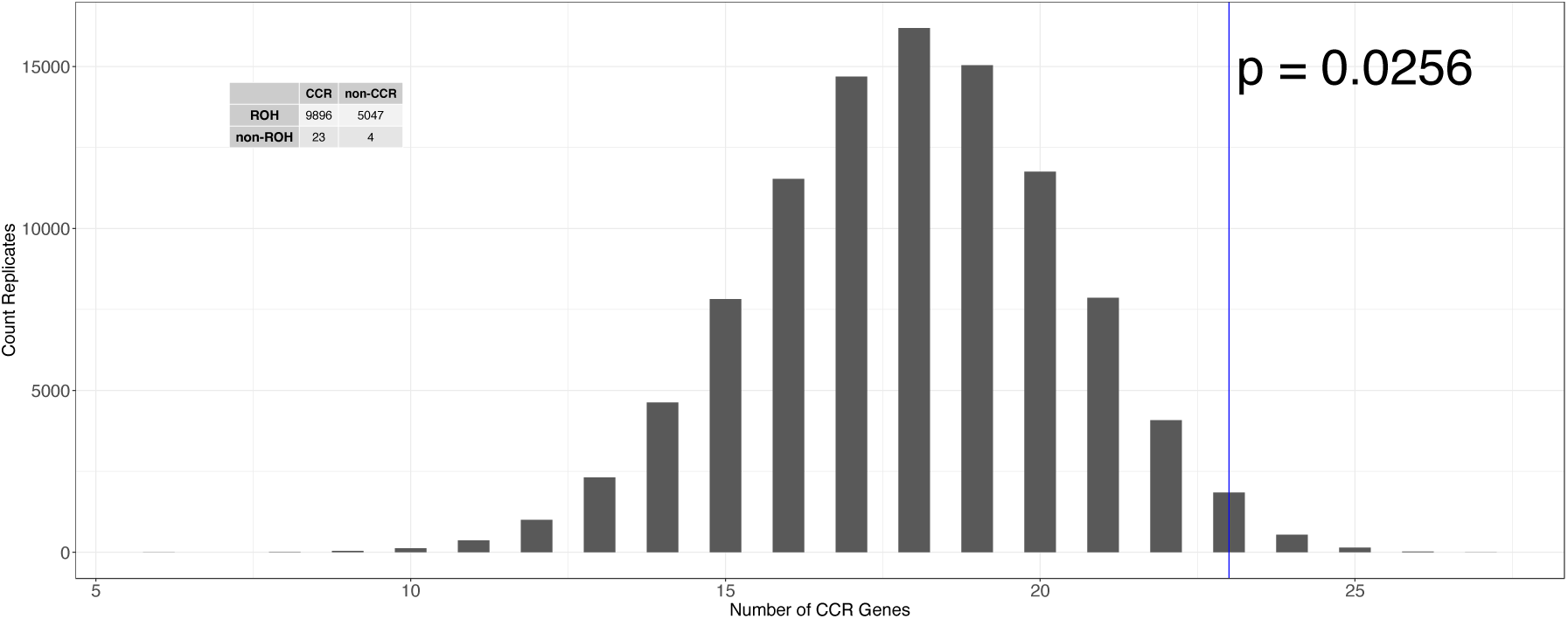
Histogram of the expected number of genes that fall into the top 10% constrained coding regions (CCRs) over 100,000 randomly drawn sets of 27 genes. The empirical data is demarcated by the blue line (p = 0.025). The contingency table shows the count of genes classified as either ROH or non-ROH and CCR or non-CCR. We observed a 2.94-fold enrichment of genes with at least one exon without an ROH, non-ROH genes, in CCRs (p = 0.041) relative to genes with an ROH.

We also tested whether our ROH analyses could be affected by low single nucleotide polymorphism (SNP) density, since the analyses thus far used only SNP genotype data. Because whole-genome sequence data would have an increased density of SNPs, we repeated our analyses using two sets of sequence data. The first dataset represents samples from four different breeds of dog: Pug (N=15) with approximately 47X coverage (47), Labrador Retriever (N=10) with approximately 30X coverage (48), Tibetan Mastiff (N=9) with approximately 15X coverage (49), Border Collie (N=7) with approximately 24X coverage (48). The second dataset was previously published (see 16), and contains 220 samples from human populations. We find that of the 27 genes with at least one exon not overlapping an ROH in any dog, three genes *ANKH, FYTTD1*, and *PRMT2* have exons not overlapping an ROH in any of the three data sets (Table S2). One of these genes, *ANKH*, has known Mendelian phenotypes that have been reported in Online Mendelian Inheritance in Man (OMIM) and is also a 95^th^ percentile CCR (50). These three genes *ANKH, FYTTD1*, and *PRMT2* all reside towards the end of the chromosome in dogs (Figure S7). Nevertheless, the relative distribution of ROHs and these three genes not containing ROH were concordant across both VCFTools and PLINK (Figure S6).

## Discussion

Here we show how the population history of dogs has increased the number of regions of the genome carried in ROHs and IBD segments, affecting phenotypes and fitness. Our work contributes to a burgeoning number of studies associating ROH burden with complex traits and is one of the first to directly show this association in dogs (3, 12, 51). Dogs provide an excellent model to examine the connection between ROH burden and disease due to their unique demographic history, as well as their simplified trait architecture due to artificial selection. Further, dogs avoid many of the challenges encountered when using human data (need for large sample size, correcting for confounding due to socio-economic status, religion, educational attainment etc.) (3, 15, 52). For example, religion has been shown to confound associations with ROH burden and major depressive disorder (53) and small sample size was shown to be a potential reason for replication failures when examining the association of ROH burden and schizophrenia (29).

Dog breeds were initially formed through domestication of one or more ancestral wild populations, in a process involving population bottlenecks. Then, over the last two hundred years or so, modern dog breeds were formed (6, 8, 10, 15, 34, 54–56). This breed formation process resulted in additional population bottlenecks. Our results confirm that the severity of the breed formation bottleneck varies (Figure 1), which to our knowledge has only been examined one other time using genetic variation data (57). Further our results match historical records quite well. For example, the Rottweiler appears to have experienced one of the most severe bottlenecks (Figure 1), and records document the Rottweiler almost disappearing in the early 1900’s and subsequently being revived by a handful of individuals (58). Conversely in breeds with a less severe bottleneck shown by the IBDNe analysis, such as the American cocker spaniel, there is a corresponding breed history that does not include a population crash, instead noting long-term breed popularity (55). Lastly, it is quite apparent that village dogs and wolves did not experience the domestication bottleneck, which we expected since they were not domesticated (9). Instead, we see much weaker recent bottlenecks in the wolves, which are likely connected to anthropogenic factors such as hunting and habitat fragmentation. While estimates of current N_e_ may be inaccurate due to low sample size, the shape trajectory of N_e_ has been shown to be more robust to the low sample size (13). Thus, our work demonstrates how recent demographic history has affected patterns of IBD and ROH within modern dog breeds.

We hypothesized that the increase in IBD and ROH in certain dog breeds would have led to an increase in the presence of recessive Mendelian diseases. Thus, we expected to observe a positive correlation between the IBD and ROH carried within a breed and the number of causal variants identified in OMIA, due to the increased probability of revealing fully or partially recessive mutations due to excess homozygosity. Instead, we found a negative association between IBD and ROH scores and the number of causal variants identified (Figure 3). Interestingly, many of the breeds with the largest amounts of IBD and ROH have had no causal variants identified through 2020. Instead, we found a positive correlation with breed popularity and the number of causal variants identified. This counterintuitive result could be caused by: 1) increased numbers of popular breed dogs seen in veterinary offices, 2) increased funding and genomic studies of disease in popular breeds (through clubs or direct-to-consumer genomics), or 3) a combination of both.

Given this ascertainment bias that we have observed in OMIA, researchers may want to shift their focus to some of these understudied breeds, as there may be more potential to discover new disease-associated variants. Ascertainment bias is not unique to OMIA and has been observed in human databases like OMIM (59). In the case of human data, authors found that OMIM contains an enrichment of diseases caused by high frequency recessive-alleles. They suggested that the bias is caused by the method through which these variants have been identified. Many variants have been identified in isolated human populations, where there may be elevated levels of relatedness, which increases the probability of mapping higher frequency deleterious variants (59).

We find that, for some traits, increased homozygosity of low frequency variants can impact phenotype. Low-frequency variants harbored in ROH likely become more important in the context of inbreeding and can lead to severe inbreeding depression. Because purebred dogs experienced severe inbreeding, in a large sample of dogs (>4,000 individuals), ROHs are expected to cover the entire genome in at least one individual (Figure S6). By searching for regions of the genome devoid of ROHs across all individuals, we can potentially identify genes containing strongly deleterious mutations, possibly underlying inbreeding depression. These regions could be lacking ROH because strongly deleterious recessive mutations lurk as heterozygotes in the founders of the breed. Then, individuals that are homozygous for these regions are no longer viable and are not sampled in our study. Similar to what has been shown in Scandinavian wolves (21), we find that there are multiple exons spanning 27 different genes where there are no ROH across all 4,300 dogs. Further, we observe that 23 of these genes were within the top 90^th^ percentile of human exons most devoid of polymorphism (the CCRs). Overall, CCRs tend to be enriched for pathogenic variants linked to clinical phenotypes (46). However, there are some CCRs that do not have any known pathogenic or likely pathogenic variants suggesting mutations in these exons could cause extreme developmental disorders or potentially be embryonic lethal (46). Therefore, studying mutations in these exons without ROH that overlap the CCR data could be fruitful both for identifying new disease phenotypes and for identifying variants with large fitness effects that could be linked to inbreeding depression or embryonic lethality.

We examined the locations of the exons without ROHs. We find that ROH tend to not occur at all within the exons of some genes or occur at exons toward the end of the gene (Figure S8). Perhaps, the location of ROH depletions could be related to nonsense-mediated decay. For example, deleterious variants within these ROH depleted regions could have large effects on gene expression, through nonsense-mediated decay, if they were to become homozygous.

Previous work has suggested there is a connection between nonsense-mediated decay and large-effect low frequency variants that disrupt splicing and potentially alter mRNA stability (60– 62). Further, nonsense-mediated decay has been shown to play a role in disease through numerous mechanisms such as patterns of inheritance, modulating the disease phenotype, and causing different traits to manifest from mutations in the same gene (63).

Our findings have implications for understanding the architecture of complex traits in other species, such as humans. Specifically, the fact that we find a relationship between ROH and certain phenotypes (Figure 2), suggests that recessive mutations play a role in some traits. Much of the existing genome-wide association studies (GWAS) in humans have largely suggested that complex traits are highly polygenic with many additive effects (64–67). These differences across species likely reflect differences in genetic architecture driven by the demographic history of the populations combined with natural selection. Nevertheless, searching for recessive variants underlying complex traits in humans may be a fruitful avenue of research. Further, variation in the amount of the genomes in ROHs across human populations (13, 19, 22, 45), could lead to population-specific architectures for complex traits. For example, causal variants in populations with a higher burden of ROHs may be more recessive and less polygenic than in populations with fewer ROHs. Future research will be needed to fully elucidate which mutations are directly responsible for severe inbreeding depression and the functional impact of these deleterious mutations. Additional work could examine which models of trait architecture (e.g. the degree of dominance and mutational target size), and demography could generate the association with ROH burden that we detected. In conclusion, the joint analysis of IBD and ROH provides considerable information about both demography and selection in the genome. This information is especially valuable in the context of fitness and disease and allows us to shed light on recent population history.

## Materials and Methods

### Genomic data

Genotype data were aggregated from two published studies (14, 15), and all original data files are publicly available through Dryad. The Fitak et al. data (14) was lifted over to CanFam3.1 then merged with Hayward et al. data (11) using PLINK (68). SNPRelate (69) was used to perform PCA (Figure 1), identify duplicate individuals, and unrelated individuals. Duplicate individuals and potential hybrid individuals were removed from the data set, and the final data set contained 4,741 breed dogs and 379 wolves. Code for merging data is available at https://github.com/jaam92/DogProject_Jaz/tree/master/MergeFitkakAndCornell.

### American Kennel Club (AKC) data

We used AKC registration data from 1926 to 2005 to compute breed popularity. This data was curated from previous work (70) and contains information for approximately 150 recognized breeds. To compute popularity through time, we drop the first entry for each breed, as this number reflects older dogs and new litters, then use the remaining data as the total number of new registrants per year. The popularity score is defined as the integral from the second entry through 2005 (available on <u>GitHub</u>).

### Online Mendelian Inheritance in Animals (OMIA) data

We downloaded all “Likely Causal” variants listed on OMIA. However, only those associated with disease, rather than other phenotypes, were used. The Likely Causal criteria is met if there at least one publication to be listed where the variant is associated with a disorder. If a variant had been identified in multiple breeds, it was counted in each breed. The total number of causal variants per breed downloaded from OMIA and is available here: https://github.com/jaam92/DogProject_Jaz/tree/master/LocalRscripts/OMIA.

### Calling identity-by-descent (IBD) segments and inferring effective population size

To call IBD segments we used software IBDSeq (35) on its default settings. The IBD segments were input into IBDNe (41) to infer effective population size. A pedigree-based recombination map (71) was used as the input genetic map for IBDNe. We set the minimum IBD segment length to 4cM, as that is the suggested length to reliably call IBD segments in genotype data when using IBDSeq. We only inferred effective population sizes in populations with at least 50 unrelated individuals and assumed a generation time of 3 years per generation for visualizing results. We use at least 50 unrelated individuals because previous work has shown that demographic trajectories are robust to smaller sample sizes, though accurate estimates of effective population size (N_e_) remain limited (13).

### Calling runs of homozygosity (ROH)

VCFTools, which implements the procedure from Auton et al. 2009 (72), was used to call ROH in all individuals. Next, we performed quality control of the raw ROH. We only kept ROHs that contained at least 50 SNPs, were at least 100 kb long, and where SNP coverage was within one standard deviation of mean SNP coverage across all remaining ROH. A file that contains the final ROHs and scripts for running quality control can be found here: https://github.com/jaam92/DogProject_Jaz/tree/master/LocalRscripts/ROH. We also called ROH using PLINK, with the following parameters: --homozyg-window-het 0 -- homozyg-snp 41 --homozyg-window-snp 41 --homozyg-window-missing 0 --homozyg-window-threshold 0.05 --homozyg-kb 500 --homozyg-density 5000 --homozyg-gap 1000 and then repeated the filtering listed above for the VCFTools ROH.

### Computing IBD and ROH Scores

We computed each population’s IBD and ROH scores using the using an approach similar to Nataksuka et al. (73). A population’s IBD score was calculated by computing the total length of all IBD segments between 4 and 20 cM and normalizing by the sample size. A population’s ROH score was computed using all ROH that passed quality control and normalizing by the sample size.

### Association test and Effect Size estimates

For these analyses, we only used the subset of breed dog data from Hayward et al. where we had phenotype information (15). We computed the association between F_ROH_ and each trait using a generalized linear mixed model which is implemented in the R package GMMAT (74). Following the protocol from Hayward et al., we did not include covariates in the association test, and included the kinship matrix as a random effect in the model, to control for population stratification due to co-variation of the amount of ROH per breed with the incidence of the phenotype in the breed. P-values were determined using a Wald test with a significance threshold of p = 0.05. We note that there is no correction for multiple testing. For more details on clinical trait ascertainment see (15). We generated the kinship matrix two different ways: 1) using the R package PC-Relate (75) on the SNP genotype matrix, and 2) by computing the total amount of the genome within a ROH that is shared between two individuals, shared ROH (SR):

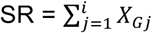

Here *X*_*Gj*_ ∈ {0, 1} and equals 1 if the genotype (*G*) at the j^th^ site falls within a ROH shared by both individuals, and equals 0 otherwise. SR was computed for each pair of individuals and bound between 0 (no sharing) and 1 (complete sharing with oneself) as follows:

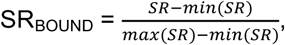

where the max amount of sharing in the equation above is the total length of the autosome and minimum is smallest SR value base pairs. Results reported in the main text are for the bounded values of SR. We also compared these results to those when not using a kinship or ROH matrix (Figure S9).

### Identifying depletions of ROH

To find the number of genes expected to contain at least one exon without a ROH, the ROHs in each individual were permuted to a new location on the same chromosome using BEDTools shuffle. Next, we created a bed file containing the permuted ROH locations and intersected this file with the exon locations from CanFam3.1 and counted the number of genes with at least one exon where we did not observe any overlap with an ROH. To control for edge effects along the chromosomes, we concatenated all 38 chromosomes into a single chromosome with a total length equivalent to the sum of all the autosome lengths. As such, the shuffled locations of ROH could occur at the ends of chromosomes. We repeated our permutation test 10,000 times to create a null distribution. The p-value was computed as the proportion of permuted datasets with as many or more genes with an exon not overlapping an ROH relative to what was seen empirically (27 genes).

To examine the overlap between genes lacking ROH and constrained coding regions (CCRs) from Havrilla et al. (46), we used BEDTools (76) to intersect non-ROHs with the top 10% (90^th^ percentile) of CCRs and exon ranges for CanFam3.1 (55) which came from Ensembl (77). Then, we tabulated the total number of genes where there was at least one exon where we never observed any overlap (including partial overlap) with a ROH (non-ROH genes) and the converse (ROH genes), as well as the count of whether these non-ROH and ROH genes fell within a CCR. Significance of the ratio of non-ROH genes relative to ROH genes within a CCR was assessed using a Fisher’s Exact test. We also computed the expected number of non-ROH genes within the 90^th^ percentile CCRs by randomly sampling an equal number of genes from the entire gene set and intersecting the randomly sampled genes with CCRs. We repeated this random sampling of genes 100,000 times to build the null distribution and computed a p-value as the proportion of Msets of the 27 genes that had at least 23 genes containing an exon in the 90^th^ percentile of CCR genes.

## Acknowledgments

We thank Eduardo Amorim, Arun Durvasula, Nelson Freimer, Jesse Garcia, Jacqueline Robinson, and Janet Sinsheimer for helpful discussions about data analysis and curation and Bob Wayne for comments on the manuscript. This material is based upon work supported by the National Science Foundation Graduate Research Fellowship under Grant DGE-1650604 awarded to JAM as well as NIH grant R35GM119856 awarded to KEL.

