## Supplementary Information for "The impact of identity-by-descent on fitness and disease in natural and domesticated *Canid* populations"

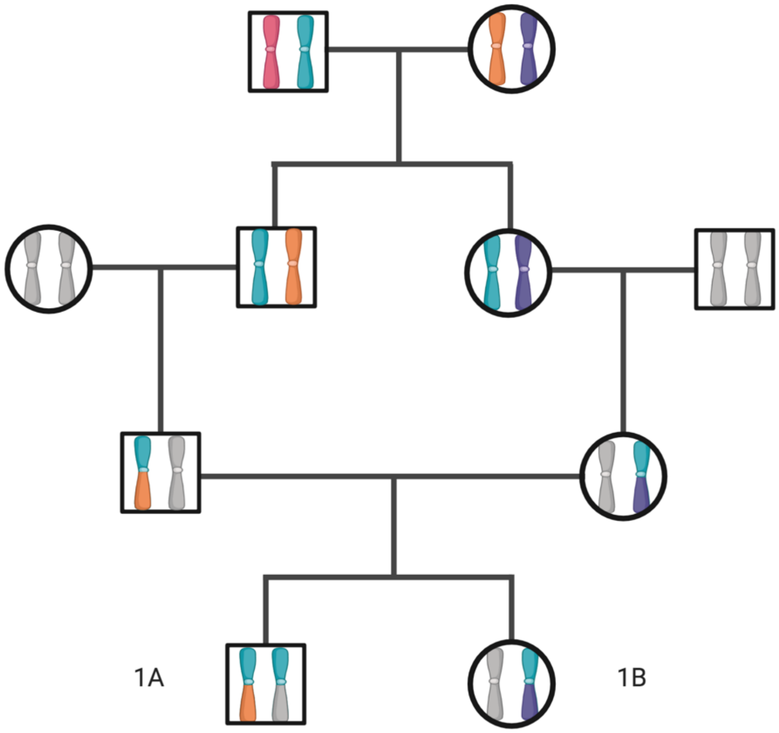


**Figure S1** Formation of runs of homozygosity (ROH) and shared identity-by-descent (IBD). This pedigree shows how runs of homozygosity (1A) and shared identity-by-descent can occur (1B), via the aqua founder chromosome. Unrelated individual’s chromosomes are shown in gray. Individual 1A inherited the aqua portion of their chromosomes identical-by-descent from both parents, which results in a run of homozygosity. Individual 1B inherited the same aqua founder haplotype as Individual 1A and thus shares an identity-by-descent segment with her brother.

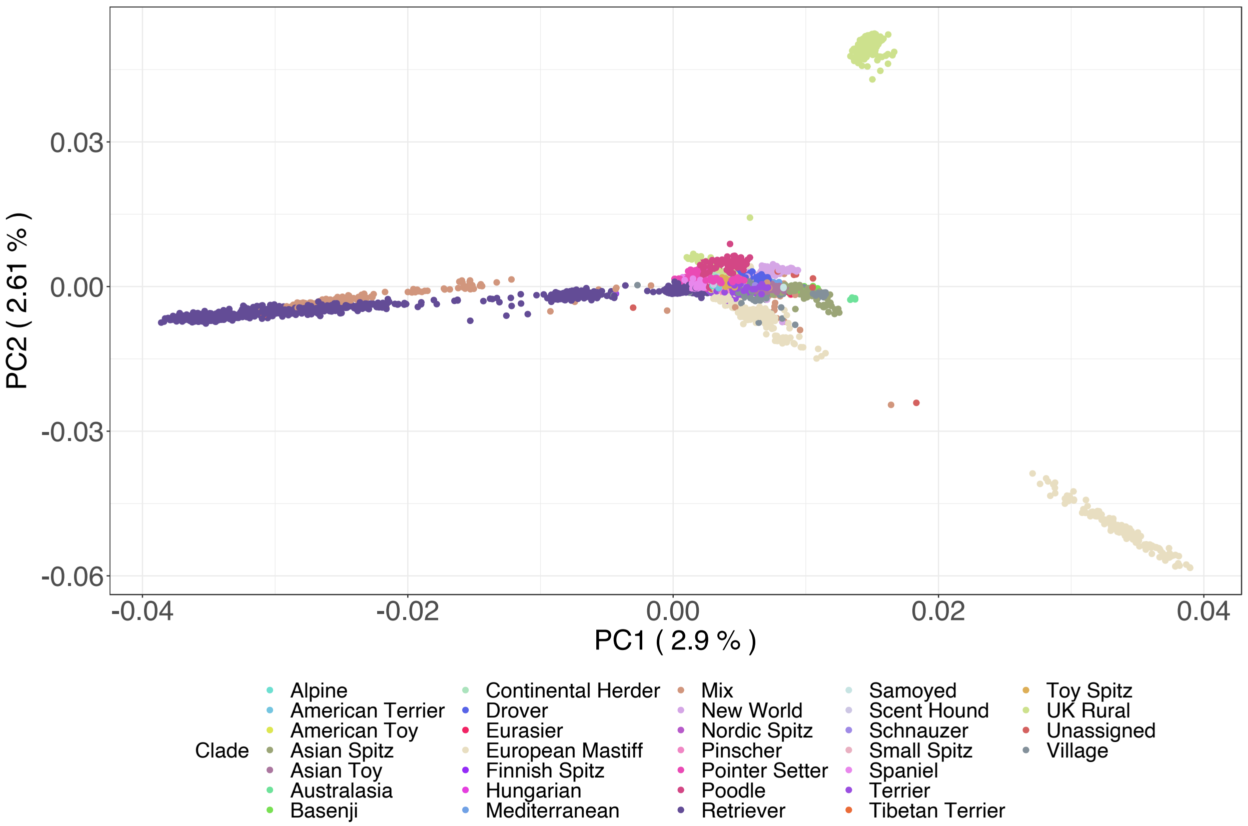


**Figure S2** PCA of dogs included in our study colored by clade. Clades are composed of multiple breeds and assignments come from two previous studies (7, 30). Dogs from a given breed tend to cluster into a single clade. There are a handful of individuals that remain unassigned which represent dogs sampled from different countries (for example a German Shepherd dog from Germany) or more divergent breeds like the Czechoslovakian Wolf Dog.

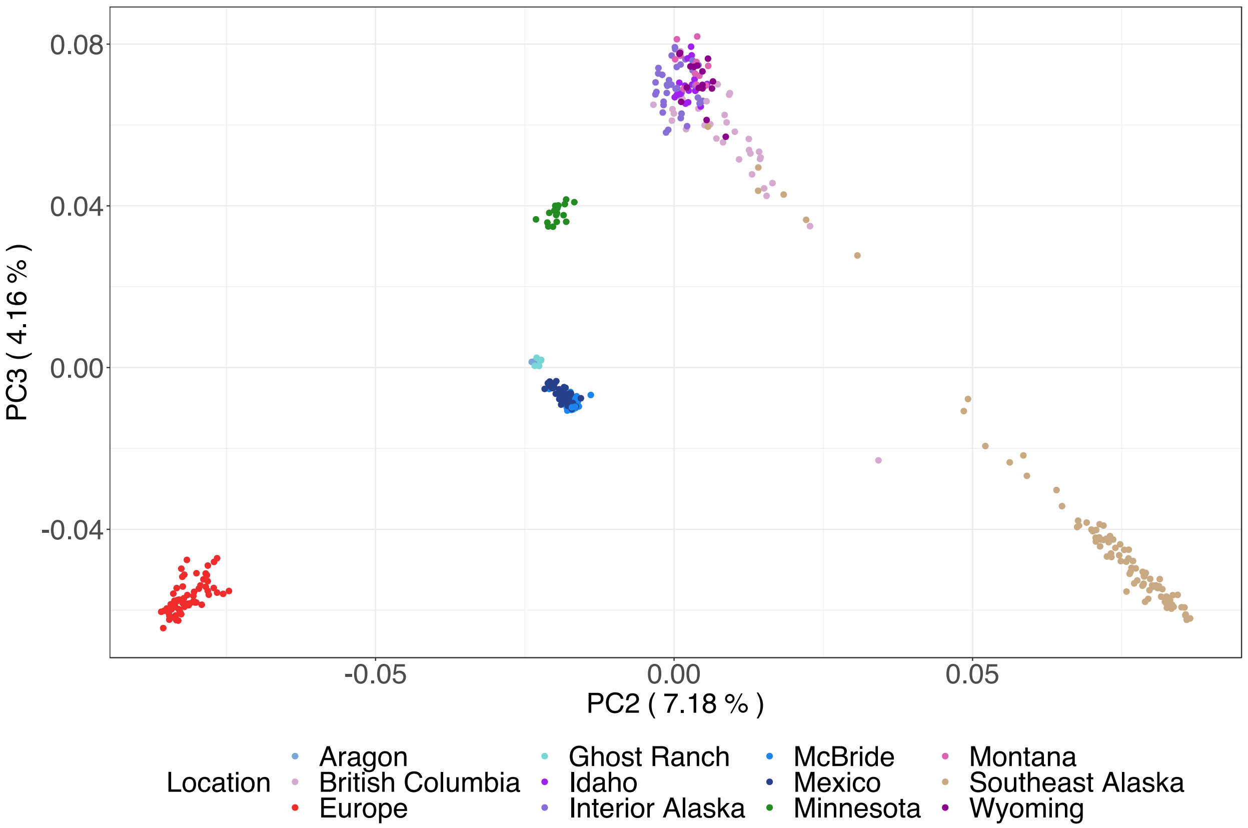


**Figure S3** PCA of wolves sampled colored by sampling location. Wolves separate by sampling location on PC2 versus PC3, whereas PC1 vs PC2 (not shown) is largely driven by continental location (America versus Europe). Wolves cluster primarily based on geographic proximity of populations.

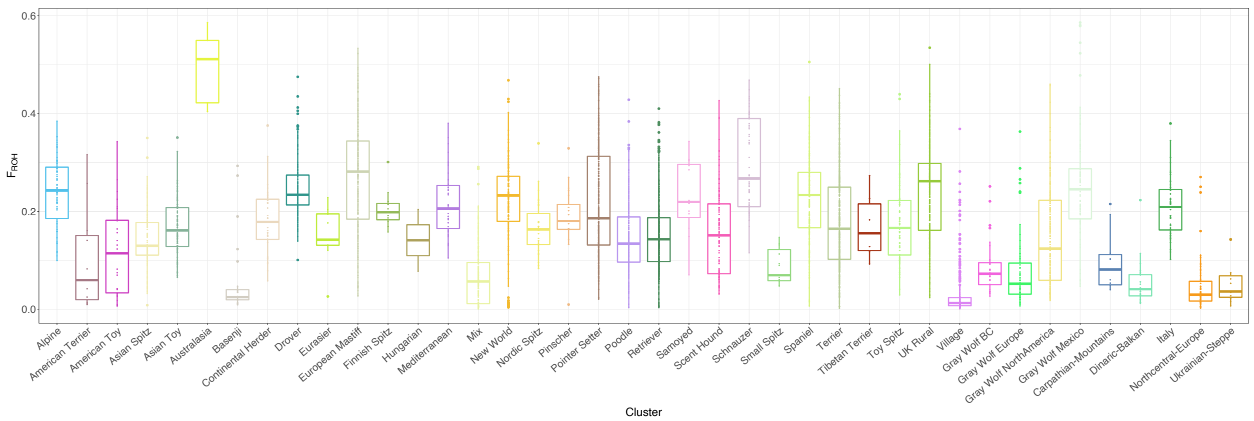


**Figure S4** $F_{ROH}$ per clade of dog or cluster of wolves. On average, dogs tend of have larger values of $F_{ROH}$ than wolves or village dogs (far right). Wolf populations with large values of $F_{ROH}$ (Mexican wolves and Italian) are known to be inbred.

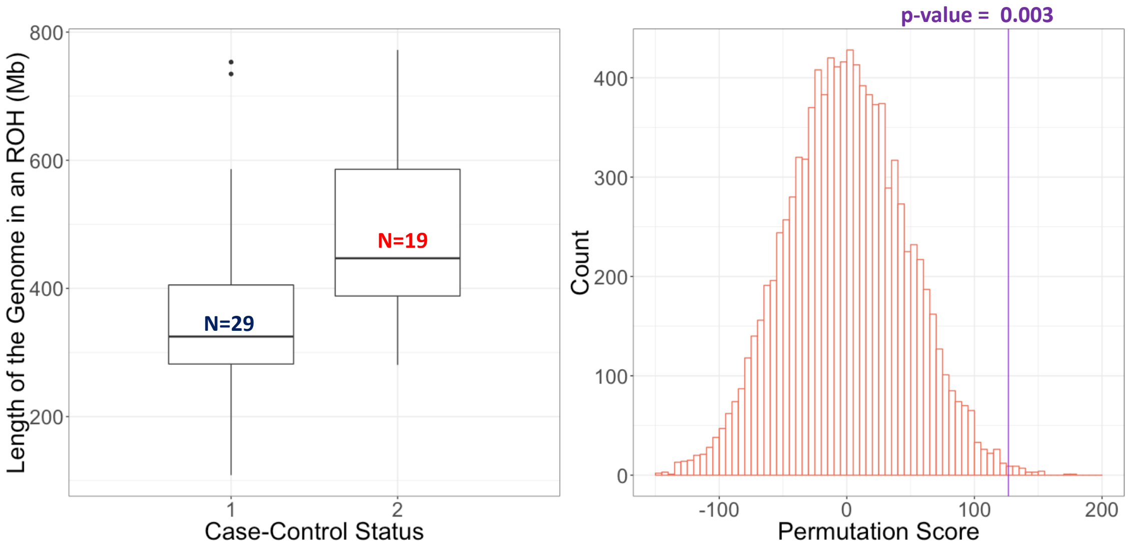


**Figure S5** Amount of the genome within an ROH in Golden Retrievers with or without lymphoma. Controls (N=29) are labelled as 1 and cases (N=19) are labelled as 2 in the graph on the left. Cases tend to have about 1.3x more of their genome within a ROH relative to controls (p-value = 0.003). Enrichment p-value was computed using a permutation test, shown on the right.

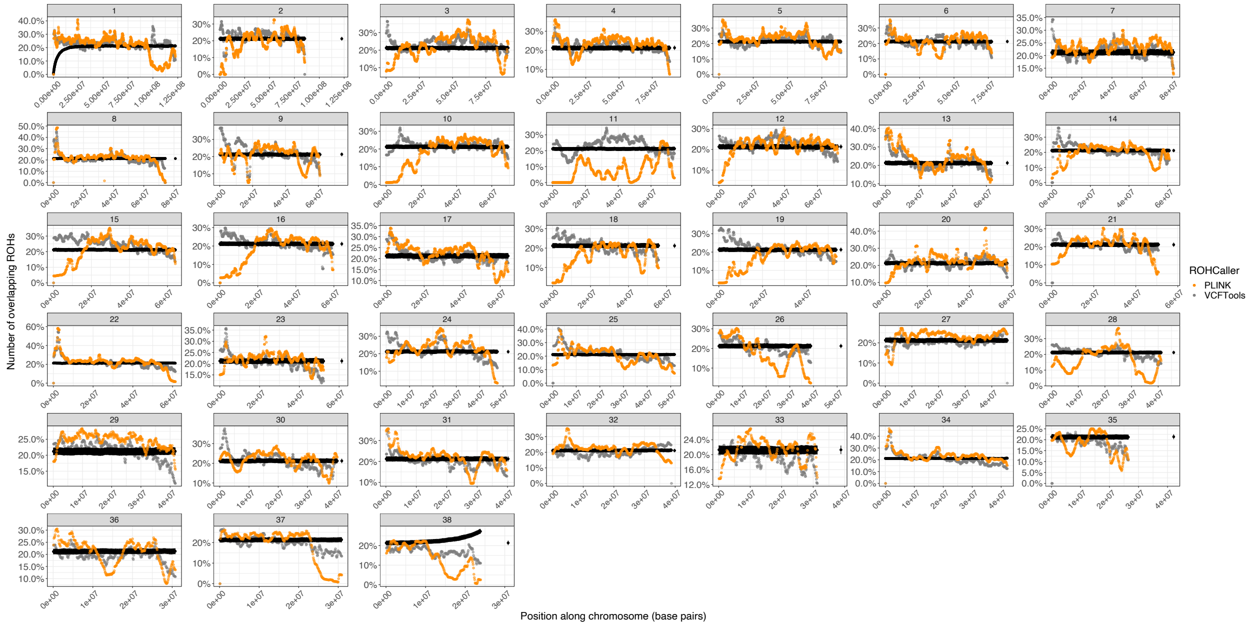


**Figure S6** Density of ROH across the genome. Here we have plotted the proportion of individuals with an ROH within 100Kb non-overlapping windows across each chromosome. Black shows permutation expectation data averaged across 10000 simulations with 95% confidence intervals, while orange (PLINK) and gray (VCFTools) show empirical ROH distribution per chromosome.

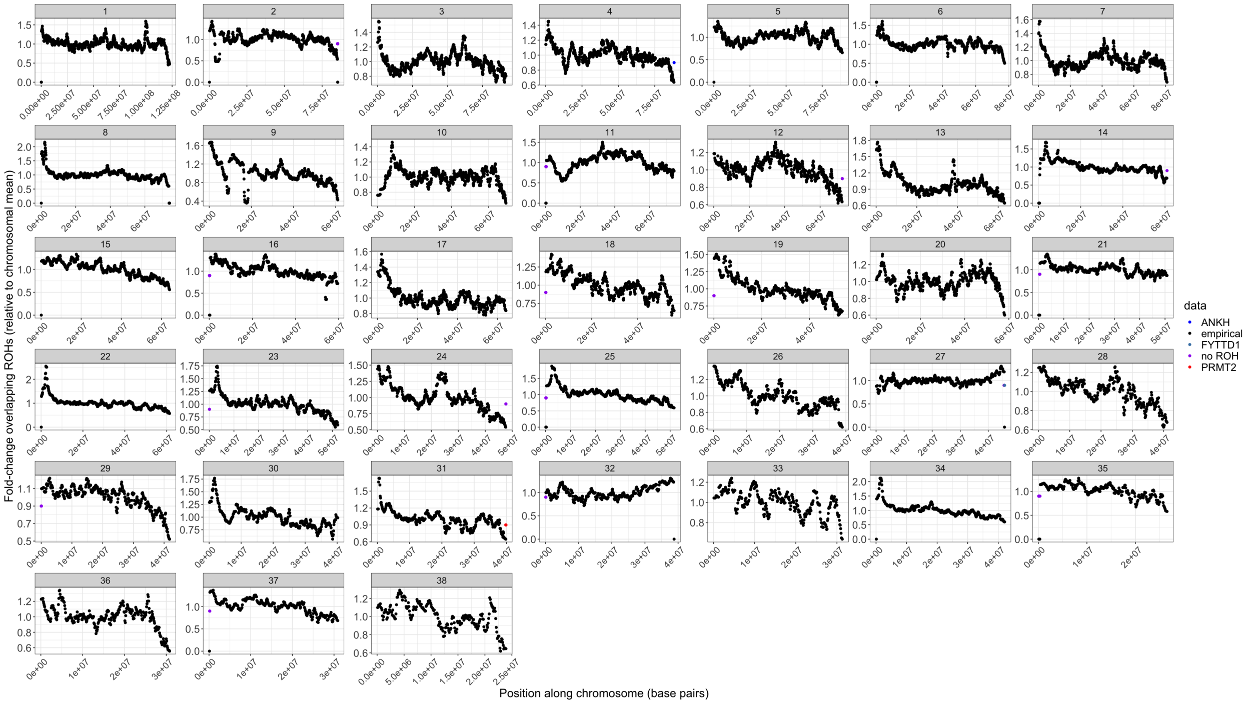


**Figure S7** Fold-change of ROH across the genome. Here we have plotted the fold-change with an ROH (using VCFTools) within 100Kb non-overlapping windows across each chromosome. We have labelled the all the exons without ROH in purple as well as the three genes that do not show a ROH in any of the three datasets (Table S2).

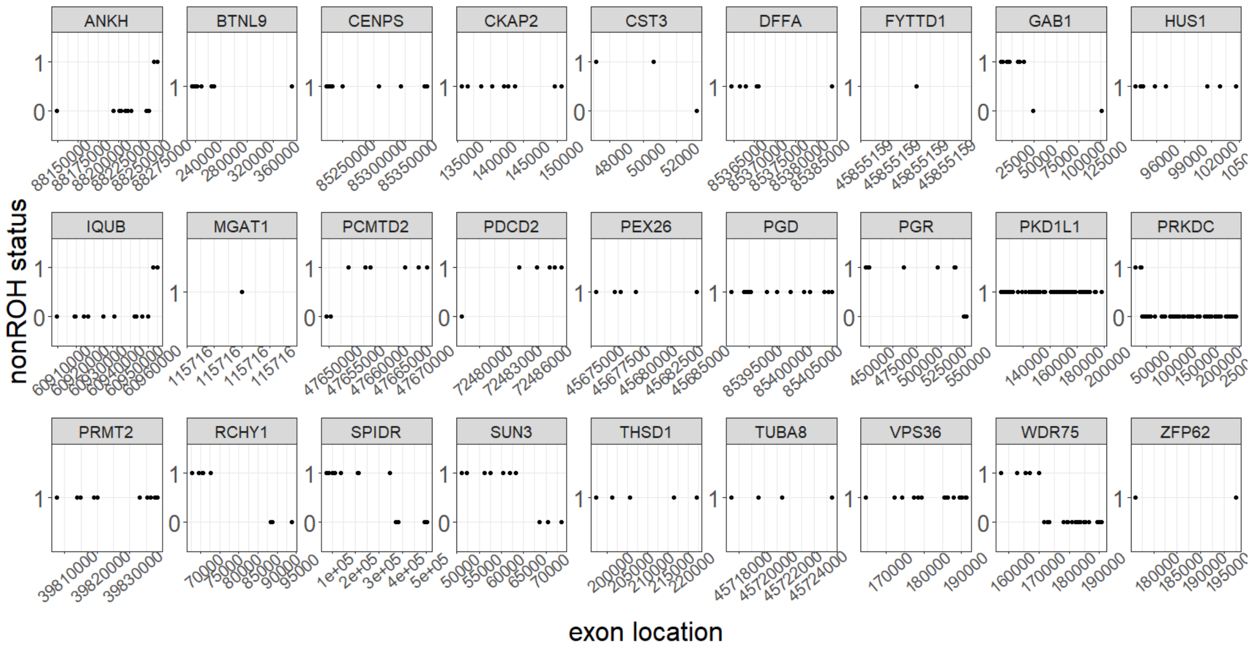


**Figure S8** Location of exons without ROH within each gene. Data above comes from nonROH regions identified in the genome after calling ROH using VCFTools. A nonROH status of one means there are no ROH overlapping that exon. A nonROH status of 0 means at least one ROH overlaps the exon. Each dot represents an exon. Overlaps can be partial. Some genes do not overlap ROH at all while others have a concentration of ROH at either the end.

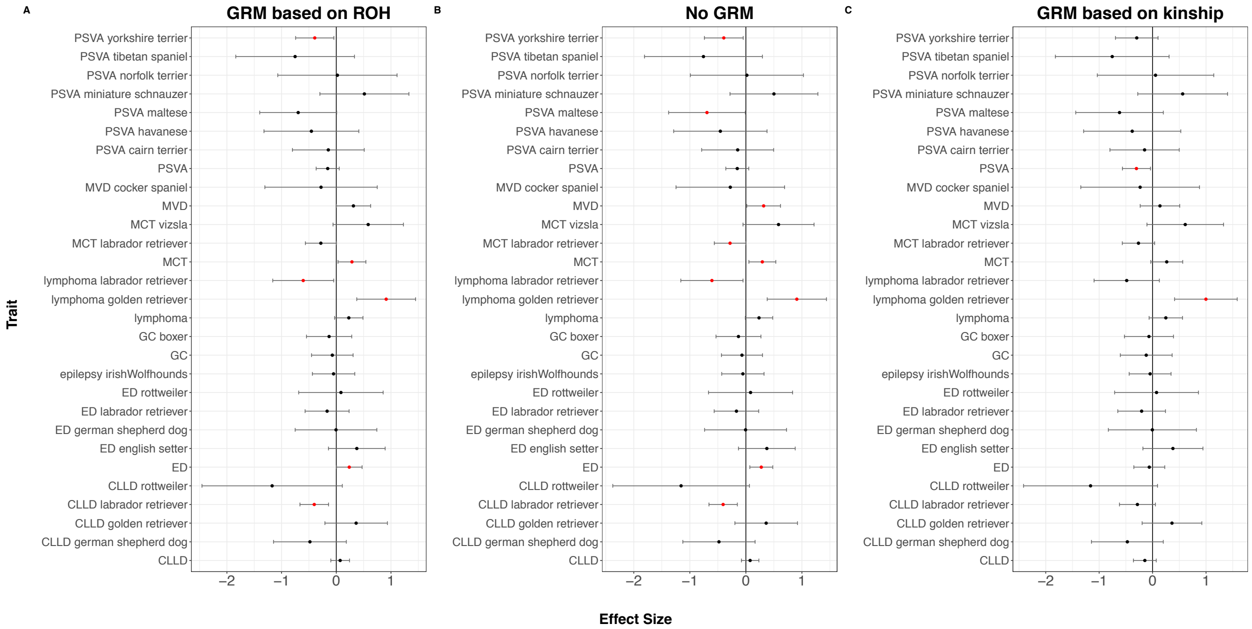


**Figure S9** Association of ROH burden with traits using different approaches to correct for breed structure. **A)** Our method (SR) which constructs a relatedness matrix based proportion of shared ROHs (the same as the results shown in Figure 2 of the main text), **B)** no correction for breed structure, and **C)** a standard correction using a kinship matrix based on genotype sharing between individuals.

| Trait | Number of controls | Number of cases | Odds-Ratio | Log(β) | Standardized Log(β) | Upper bound | Lower bound | P |
|---|---|---|---|---|---|---|---|---|
| CLLD German Shepherd dog | 24 | 24 | 0.618 | -0.482 | -1.420 | 0.183 | -1.147 | 0.156 |
| CLLD Golden Retriever | 32 | 28 | 1.438 | 0.363 | 1.247 | 0.934 | -0.208 | 0.213 |
| **CLLD Labrador Retriever** | **173** | **114** | **0.669** | **-0.403** | **-3.015** | **-0.141** | **-0.664** | **0.003** |
| CLLD Rottweiler | 11 | 11 | 0.309 | -1.173 | -1.792 | 0.110 | -2.456 | 0.073 |
| CLLD | 399 | 271 | 1.074 | 0.071 | 0.814 | 0.242 | -0.100 | 0.416 |
| ED English Setter | 52 | 27 | 1.456 | 0.375 | 1.419 | 0.894 | -0.143 | 0.156 |
| ED German Shepherd dog | 45 | 10 | 0.995 | -0.005 | -0.013 | 0.741 | -0.751 | 0.989 |
| ED Labrador Retriever | 180 | 30 | 0.846 | -0.167 | -0.816 | 0.235 | -0.570 | 0.414 |
| ED Rottweiler | 30 | 11 | 1.089 | 0.085 | 0.216 | 0.857 | -0.687 | 0.829 |
| **ED** | **633** | **113** | **1.268** | **0.238** | **1.983** | **0.473** | **0.003** | **0.047** |
| Epilepsy Irish Wolfhounds | 167 | 34 | 0.952 | -0.050 | -0.251 | 0.338 | -0.437 | 0.802 |
| GC Boxer | 74 | 40 | 0.877 | -0.131 | -0.621 | 0.283 | -0.545 | 0.535 |
| GC | 91 | 46 | 0.930 | -0.072 | -0.372 | 0.308 | -0.452 | 0.710 |
| **Lymphoma Golden Retriever** | **48** | **43** | **2.491** | **0.913** | **3.324** | **1.451** | **0.374** | **0.001** |
| **Lymphoma Labrador Retriever** | **62** | **22** | **0.546** | **-0.604** | **-2.125** | **-0.047** | **-1.162** | **0.034** |
| Lymphoma | 138 | 138 | 1.257 | 0.229 | 1.734 | 0.488 | -0.030 | 0.083 |
| MCT Labrador Retriever | 107 | 107 | 0.754 | -0.282 | -1.950 | 0.001 | -0.565 | 0.051 |
| MCT Vizsla | 26 | 26 | 1.793 | 0.584 | 1.779 | 1.227 | -0.059 | 0.075 |
| **MCT** | **146** | **146** | **1.332** | **0.286** | **2.207** | **0.541** | **0.032** | **0.027** |
| MVD Cocker Spaniel | 11 | 11 | 0.757 | -0.278 | -0.531 | 0.749 | -1.306 | 0.595 |
| MVD | 95 | 95 | 1.368 | 0.313 | 1.950 | 0.629 | -0.002 | 0.051 |
| PSVA Cairn Terrier | 23 | 21 | 0.865 | -0.146 | -0.434 | 0.511 | -0.802 | 0.664 |
| PSVA Havanese | 15 | 15 | 0.635 | -0.455 | -1.028 | 0.412 | -1.322 | 0.304 |
| PSVA Maltese | 24 | 24 | 0.499 | -0.696 | -1.939 | 0.008 | -1.400 | 0.053 |
| PSVA Miniature Schnauzer | 17 | 17 | 1.672 | 0.514 | 1.238 | 1.328 | -0.300 | 0.216 |
| PSVA Norfolk Terrier | 10 | 10 | 1.021 | 0.021 | 0.037 | 1.111 | -1.070 | 0.970 |
| PSVA Tibetan Spaniel | 13 | 10 | 0.471 | -0.753 | -1.360 | 0.332 | -1.838 | 0.174 |
| **PSVA Yorkshire Terrier** | **105** | **57** | **0.675** | **-0.394** | **-2.209** | **-0.044** | **-0.743** | **0.027** |
| PSVA | 216 | 171 | 0.855 | -0.157 | -1.455 | 0.054 | -0.368 | 0.146 |

**Table S1** Association between different phenotypes (rows) and the amount of the genome in a ROH Traits are either split by breed or aggregated across multiple breeds. Bolded results are significant. Note, p-values have not been corrected for multiple testing. Phenotype abbreviations: cranial cruciate ligament disease (CCLD); elbow dysplasia (ED); granulomatous colitis (GC); mast cell tumor (MCT); mitral valve degeneration (MVD); and portosystemic vascular anomalies (PSVA).

| Organism | Data Type | Number of individuals | Number of Genes without ROH |
|---|---|---|---|
| **Dog** | Genotype | 4342 | 27 |
| **Dog** | Sequence | 41 | 124 |
| **Human** | Sequence | 220 | 4939 |
| **Shared** | N/A | N/A | 3 |

**Table S2** Genes without ROH across multiple datasets. We called ROH in three different datasets. The first dataset is the discussed in the main text. The second dataset is sequence data from dogs and the third is sequence data from humans (Mooney et al. 2018). There were 3 genes (*ANKH*, *PRMT2*, *FYTTD1*) that were found to have at least one exon without an ROH across all three datasets.
